# Transcriptome Landscape Reveals Underlying Mechanisms of Ovarian Cell Fate Differentiation and Primordial Follicle Assembly

**DOI:** 10.1101/803767

**Authors:** Jun-Jie Wang, Wei Ge, Qiu-Yue Zhai, Jing-Cai Liu, Xiao-Wen Sun, Wen-Xiang Liu, Lan Li, Chu-Zhao Lei, Paul W. Dyce, Massimo De Felici, Wei Shen

**Affiliations:** College of Life Sciences, Institute of Reproductive Sciences, Qingdao Agricultural University, Qingdao 266109, China; Key Laboratory of Animal Genetics, Breeding and Reproduction of Shaanxi Province, College of Animal Science and Technology, Northwest A&F University, Yangling 712100, China; Department of Animal Sciences, Auburn University, Auburn, AL 36849, USA; Department of Biomedicine and Prevention, University of Rome Tor Vergata, Rome 00133, Italy

**Keywords:** Primordial follicle assembly, Oocyte, Granulosa cells, Single-cell transcriptome

## Abstract

Primordial follicle assembly in mammals occurs at perinatal ages and largely determines the ovarian reserve available to support the reproductive lifespan. The primordial follicle structure is generated by a complex network of interactions between oocytes and ovarian somatic cells that remain poorly understood. In the present research, using single-cell RNA sequencing performed over a time-series on mouse ovaries coupled with several bioinformatics analyses, the complete dynamic genetic programs of germ and granulosa cells from E16.5 to PD3 are reported for the first time. The time frame of analysis comprises the breakdown of germ cell cysts and the assembly of primordial follicles. Confirming the previously reported expression of genes by germ cells and granulosa cells, our analyses identified ten distinct gene clusters associated to germ cells and eight to granulosa cells. Consequently, several new genes expressed at significant levels at each investigated stage were assigned. Building single-cell pseudo temporal trajectories five states and two branch points of fate transition for the germ cells, and three states and one branch point for the granulosa cells were revealed. Moreover, GO and ClueGO term enrichment enabled identifying biological processes, molecular functions and cellular components more represented in germ cells and granulosa cells or common to both cell types at each specific stage. Finally, by SCENIC algorithm, we were able to establish a network of regulons that can be postulated as likely candidates for sustaining germ cell specific transcription programs throughout the investigated period.

## INTRODUCTION

Gametogenesis is a finely regulated and complex process beginning from germline specification that gives rise to primordial germ cells (PGCs) and ending with mature oocytes or sperms. In female mammals, primordial follicles (PFs) are the first functional unit of reproduction, each comprising a single oocyte derived from a PGC surrounded by supporting somatic cells termed granulosa cells. At birth, the whole of PFs constitute the ovarian reserve (OR), responsible for continued folliculogenesis and oocyte maturation throughout the adult life (Kerr, Myers et al., 2013, Tingen, Kim et al., 2009). The PF population has long been believed to be non-renewable, although this notion has recently been challenged (Johnson, Canning et al., 2004, White, Woods et al., 2012). In any case, during the reproductive lifespan, the number of PFs progressively diminish due to atresia as well as recruitment, maturation, and ovulation. The depletion of the reserve, in primates, is the major determining driver of menopause in adult females and of various conditions of infertility in young individuals grouped as having premature ovarian failure (POF) (Skinner, 2005).

In the mouse, germline specification occurs before gastrulation in the epiblast approximately at embryonic day (E) 6.25, in response primarily to bone morphogenetic proteins (BMPs) signals (Hayashi, de Sousa Lopes et al., 2007, Seki, Hayashi et al., 2005). Around E7.0-8.0, PGCs are identifiable in the extraembryonic mesoderm at the angle between the allantois and the yolk sac (Lawson, Dunn et al., 1999). From here, PGCs move into the embryo proper to colonize, between E11.5-12.5, the gonadal ridges forming from the coelomic epithelium (Nef, Stévant et al., 2019, Stévant & Nef, 2019). After active proliferation, PGCs/oogonia cease dividing at E13.5 and enter meiosis to form oocytes (Chassot, Gregoire et al., 2011). The oocytes are closely associated in clusters, termed germ cell nests, in which they are connected to each other through cytoplasmic bridges formed because of incomplete cytokinesis during mitosis (Pepling & Spradling, 1998). Oocyte loss and cyst breakdown begin after birth in the cortical region of the ovary, but in the medullar region, these processes begin as early as E17.5 (Pepling, Sundman et al., 2010). Nests breakdown begins after oocytes undergo the first meiotic block at the diplotene stage of prophase I. Failure of mouse oocytes to reach the diplotene stage impairs this process and consequently PF assembly (Paredes, Garcia-Rudaz et al., 2005, Wang, Zhou et al., 2017), but it’s still reported the two events of meiotic progression and primordial follicle formation were independent (Dutta, Burks et al., 2016). As reported above, breakdown of the cystic nests is associated with massive oocyte degeneration that in the mouse has been reported to occur following programmed cell death mainly in the form of apoptosis and autophagy (Klinger, Rossi et al., 2015, Pepling, 2006, Wang et al., 2017). The surviving single oocytes become enveloped by granulosa cells surrounding the nest, thereby producing the PFs. Synchronized development of oocytes and granulosa cells, as well as their interactive communication are required for efficient nest breakdown and the subsequent PF assembly.

Both in mouse and human, mutations of *Wnt4* and *Rspo1* genes induce failure of pre-granulosa cell differentiation, impair nest breakdown, and finally result in POF (Chassot, Bradford et al., 2012). Recent studies in the mouse identified two waves of granulosa cell differentiation that contribute to two discrete populations of follicles (Zheng, Zhang et al., 2013, Zheng, Zhang et al., 2014). The precursors of both populations arise from the ovarian surface epithelium. The first, identifiable by the early expression of the transcription factor FOXL2, encircles the oocyte nests and eventually form the earliest PFs that develop within the ovary medulla and undergo rapid maturation and atresia. The second population is composed of cells expressing LGR5, a likely receptor of WNT4/RSPO1, that subsequently express FOXL2 and downregulate LGR5 and will form the OR of PFs in the cortex (Mork, Maatouk et al., 2012, Rastetter, Bernard et al., 2014). Other somatic cell transcription factors such as IRX3 and −5 and the oocyte transcription specific factors FIGLA, SOHL1 and −2, LHX8, NOBOX, TAF4b and AHR also appear to play a role in nest break down and PF assembly both in mice and humans (Grive & Freiman, 2015, Pepling, 2012). Several signalling exchanges between oocytes and pre-granulosa cells are also implicated in such processes, such as Notch, KITL/KIT system, neurotrophins, TGFβ and its family members GDF9, BMP15, ActA, Inhibin, Follistatin, anti-Mullerian hormone, as well estrogens, progesterone and finally FSH (Tingen et al., 2009, Wang et al., 2017).

Despite these results and previous transcriptome analyses performed in rat (Kezele, Ague et al., 2005, Nilsson, Zhang et al., 2013) and mouse fetal and early post-natal ovaries (Bonnet, Cabau et al., 2013, Tan, Zhang et al., 2018), the precise molecular mechanisms underlying oocyte survival/death, granulosa cell differentiation and the crosstalk among them during PF assembly are still incomplete. Moreover, all these studies were performed on entire ovary or follicles so that the contribution of each cell type to the transcriptomes was not distinguishable. The recent advent of single-cell RNA-seq (scRNA-seq) technologies make it possible to identify specific cell subpopulations and the genetic programs regulating their differentiation in reproductive organs (Stévant, Kühne et al., 2019, Stévant & Nef, 2019).

In the present paper, in order to achieve such goals mainly focusing on PF assembly in the mammalian ovary, late fetal and early post-natal mouse ovaries were subjected to scRNA-seq analyses. The results of such analyses and the bioinformatics elaboration reported here will certainly be useful to elucidate the molecular control underlying PF assembly and select candidate regulatory factors for further investigation.

## RESULTS

### scRNA-seq identified several types and subpopulations of germ cells and somatic cells in late fetal and early post-natal ovaries

To trace the distinct cell lineages and characterize the gene expression dynamics of ovarian cells during the crucial period of ovarian development lasting from late fetal and early post-natal age, ovaries were collected from E16.5, PD0 and PD3 mice and subjected to immunohistological and scRNA-seq analyses. As expected, immunohistological sections showed that the germ cell nests gradually broke down, while the percentage of germ cells enclosed in PFs increased from 9.95±1.38 % at E16.5 to 46.02±1.46 % at PD3 (Figures 1A and 1B).

**Fig. 1.**
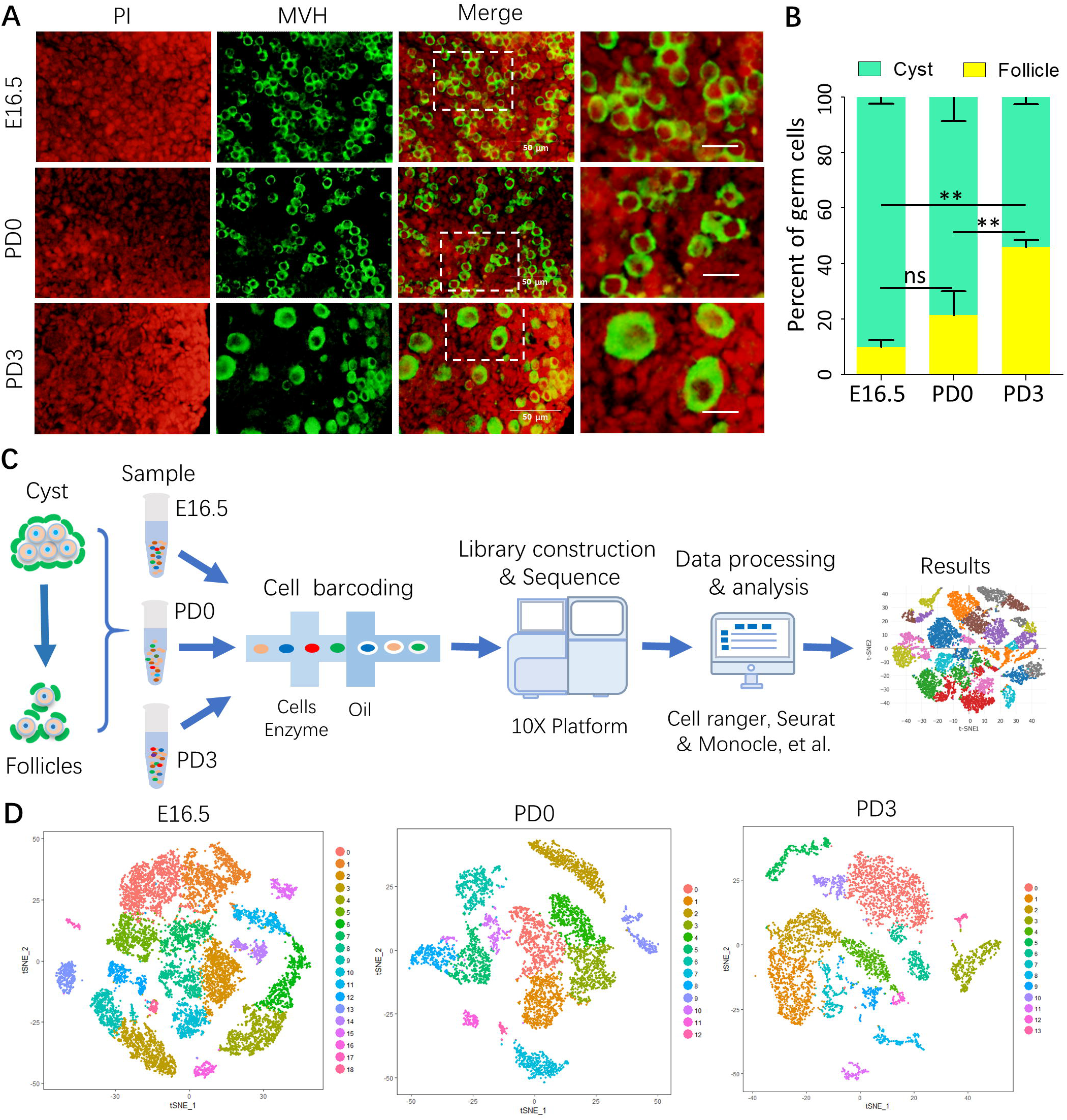
Single-cell RNA sequence analysis of ovarian cells. (A) Representative images of fetal ovaries at E16.5 and postnatal ovaries at PD0 and PD3. Germ cells were labelled with MVH (green) and nuclei counterstained with propidium iodide (PI, red). Scale bar: 50 μm. (B) Percentage of germ cells in cysts or follicles at the indicated stages. (C) Schematic diagram of the scRNA-seq analysis procedure. Ovaries were isolated and disaggregated into single cell suspensions; cells were barcoded and used for library construction onto 10x genomics platform and the data produced after sequencing analyzed by dedicated software. (D) tSNE (t-distributed stochastic neighbor embedding) plots of ovarian cell subsets at E16.5, PD0 and PD3.

Ovarian cell populations were then prepared, and barcoded following single cell capture, and RNA sequenced (Figure 1C). High-quality cells were 11,526 in E16.5, 5,466 in PD0 and 5,962 in PD3, and the mean number of genes expressed in each sample was 1,781, 2,916 and 2,832, respectively. According to t-SNE (t-distributed stochastic neighbor embedding), they were grouped in 19, 13 and 14 clusters, at E16.5, PD0 and PD3, respectively (Figure 1D). When the cell clusters of the three developmental stages were grouped, six cell types were identified (Figure 2A) that, according to the distinct genome signatures, could be subdivided as follows: germ cells distributed in six distinct clusters (3, 6, 7, 15, 16, 19), granulosa cells in six distinct clusters (1, 4, 5, 8, 12, 14), epithelial cells in three distinct clusters (0, 9, 13), erythrocytes in three distinct clusters (2, 10, 20), immune cells in two distinct clusters (17, 18) and endothelial cells only in one cluster (11) (Figures 2B and 2C). The expression of representative genes and other genes known to be specific for each cell type were reported for each cluster as dot plots in Figure 2D. In Figure 2E, the percent changes within the ovarian populations of the six identified cell types, during the analyzed developmental period, were reported. In particular, it can be observed that germ cells underwent a percent increase from E16.5 (17.8 %) to PD0 (54.4 %), and a rapid decrease at PD3 (6.8 %), while granulosa cells represented about 40% at E16.5, 20% at PD0 and again at around 40% at PD3.

**Fig. 2.**
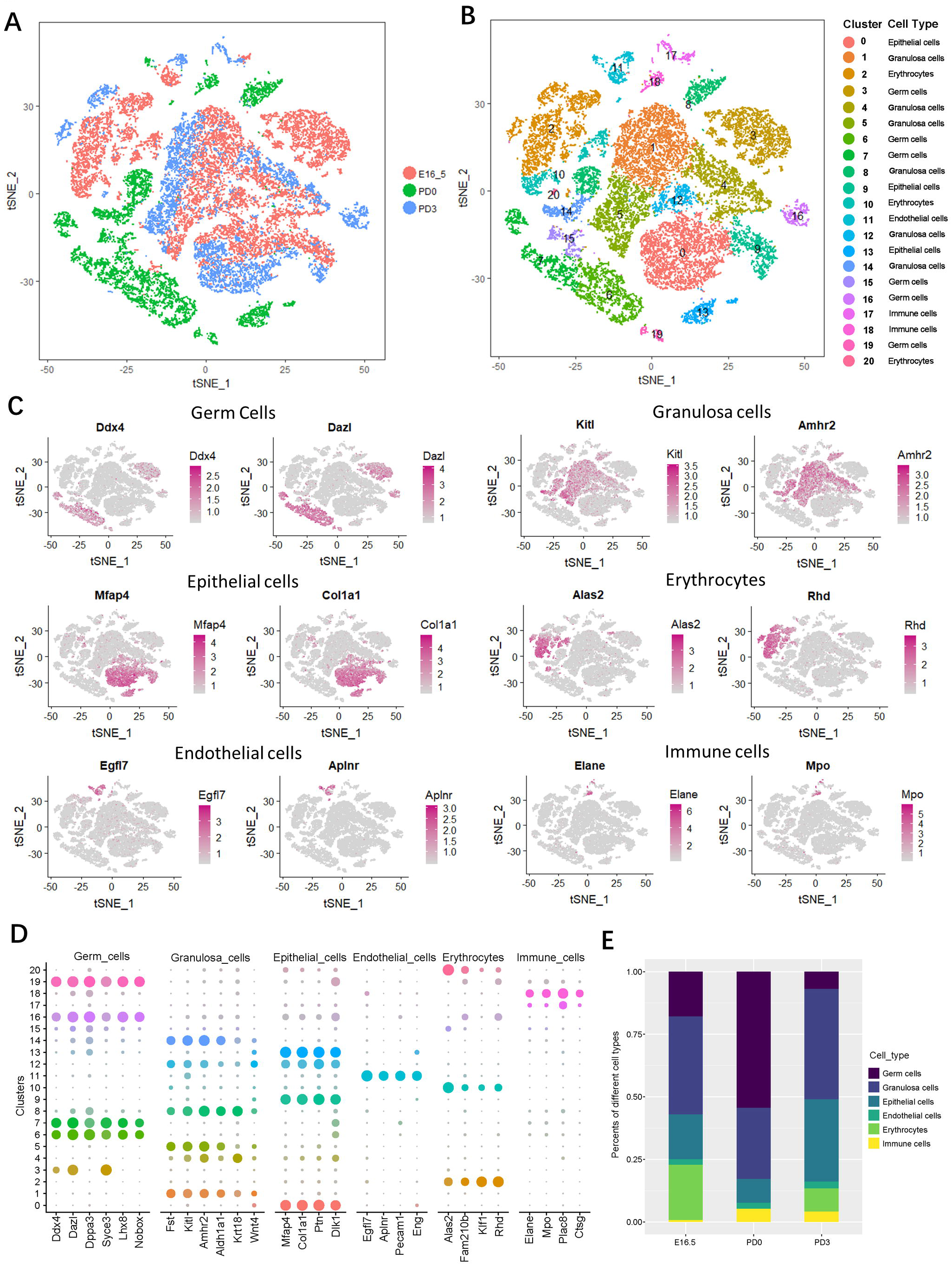
Identification of ovarian cell types by t-SNE plots. (A and B) t-SNE plots of an integrated analysis based on ovarian cells grouped by developmental stage (A) and expression patterns (B). (C) Feature plots of specific marker genes of different cell types, including germ cells, granulosa cells, epithelial cells, erythrocytes, endothelial cells and immune cells. (D) Dot plot of marker genes of the six ovarian cell types based on 20 t-SNE clusters. (E) Percentages of the six ovarian cell types at E16.5, PD0 and PD3.

The detailed identification of germ cells at each stage by *Ddx4* and *Dazl* expression is shown in Figures S1A-S1C. The canonical correlation analysis (CCA) of the expressed genes in all cell types at each developmental stage, is reported in Figures S1D and S1E. The top genes driving canonical correlation 1 (CC1) and 2 (CC2) include key genes such as *Smc1b*, *Sycp1*, *Syce2* and *Sycp3* involved in germ cell meiosis (Bolcun-Filas, Costa et al., 2007, Garcia-Cruz, Brieno et al., 2010), *Dazl* and *Xist* driving germ cell transcription (Chen, Welling et al., 2014, Sado & Sakaguchi, 2013), and *Sox4* and *Wnt6* widely expressed in folliculogenesis (Harwood, Cross et al., 2008, Zhang, Yan et al., 2018, Zhao, Arsenault et al., 2017) (Figure S1F). A heatmap of the top ten marker genes for all six cell types is displayed in Figure S1G (Supplemental Table 1).

### Genetic dynamics of female germ cells from the cyst stage to the primordial follicle formation

In the first t-SNE analysis, six germ cell populations were identified expressing genes such as *Ddx4*, *Dazl*, *Dppa3*, Syce3, *Lhx8* and *Nobox* in which cluster 3 were allocated at E16.5, clusters 6, 7, 15 and 19 to PD0 and cluster 16 to PD3 (Figures 2A-2D, and 3A). By adding to the analysis genes known to be expressed throughout the female germ cell development at the beginning of meiosis (*Stra8*, *Prdm9*, *Stat3*, *Scyp3*, *Scyp1, Rhox9*) (Bolcun-Filas & Schimenti, 2012, Gu, Tekur et al., 1998, Imai, Baudat et al., 2017, Takasaki, Rankin et al., 2001), during primordial follicle formation (*Figla*, *Sohlh1, Ybx2*) (Pepling, 2012, Wang et al., 2017, White et al., 2012), and early during oocyte growth (*Gdf9*, *Zp2* and *Zp3*) (Bayne, Kinnell et al., 2015, Dean, 2002, Rajkovic, Pangas et al., 2004, Zhang et al., 2018), ten clusters were generated in which clusters 1, 4, 5, 6 were allocated to E16.5, clusters 0, 2, 3, 8, 9 to PD0, and cluster 7 to PD3 (Figures 3B and S2A and S2B). The heat map of the five top expressed genes of each cluster is shown in Figure 3C. According to these analyses, a genetic dynamic model, including pre-, early- and late-follicular stages during germ cell development was drawn (Figures 3D and 3E). Moreover, the percentage changes of cells at these stages from E16.5 to PD3 are shown in Figure 3F. Several new genes expressed by germ cells at the pre- (i.e. *Rhox9*, *M1ap*, *Hspb1, Inca1*), early- (i.e. *Eif4a1, G3bp*, *Id1*, *Gdp1*) and late- (i.e. *Ooep*, *Xdh, Padi6*) follicle development stages were identified (Figures 3D and 3E). The expression at the protein level of HSBP11, G3BP2 and XDH at the pre-, early- and late follicular stages, respectively, were confirmed (Figures S3A, S3B and S3C).

**Fig. 3.**
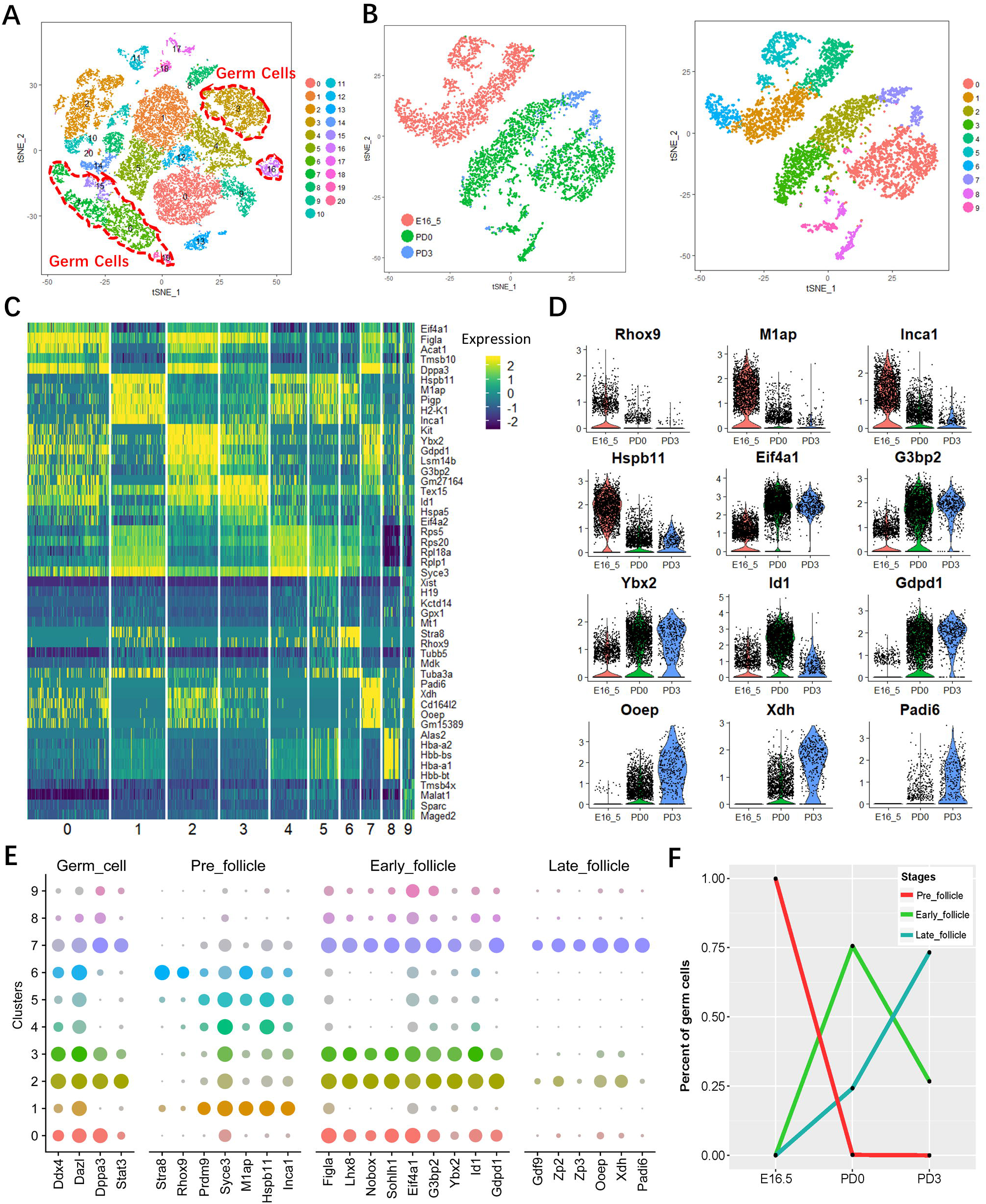
Molecular characterization of the germ cell subsets. (A) t-SNE plot of ovarian cell clusters in which germ cell subsets were marked with red dotted line. (B) Cluster analysis of germ cells with t-SNE plots based on developmental timeline (left) and transcriptional patterns (right). (C) Heatmap of top marker genes of the germ cell clusters. (D) Vlnplots of expression level of representative genes of this group as a function of developmental timeline. (E) Dot plot of general and stage-specific (Pre-, Early- and Late-follicle) germ cell marker genes expressed in the ten germ cell clusters. (F) Percent changes of Pre-, Early- and Late-follicle germ cells at E16.5, PD0 and PD3.

### Fate transition of oocytes along pseudotime trajectories

To dissect the fate determination of germ cells throughout the investigated period, they were ordered along pseudotime trajectories according to the gene expression reported above. Five states were obtained in which the majority of cells at the pre-follicle stage belonged to states 1 and 2, while those at the early and late follicle stages were in states 3, 4 and 5. In such pseudotime trajectories, two branching points were identified. Point 1 at which cyst germ cells were at state 1, considered to be derived from the less differentiated cells of state 2 and, enter state 3, and point 2, at which follicular oocytes branched to 4 and 5 states (Figures 4A and 4B and S2C). As noted, at PD0 and PD3, although the majority of cells were at states 3, 4 and 5, a small germ cell subpopulation remained at state 1 (Figure 4C). At branch point 1, four gene clusters with distinct patterns were identified likely involved in the commitment of cyst germ cells into follicular oocytes (point 1); clusters 1, 2, 3 and 4 including 2,014, 2,087, 2,406 and 911 genes, respectively. Figure 4D shows the heatmap of the dynamics of gene expression in the germ cells at cyst and follicle stages at branch point 1. Cluster 1 includes genes with low steady-state expression in germ cell cysts and highly up-regulated in follicular oocytes, cluster 2 genes down-regulated in germ cells cysts and with moderate increased expression in follicular oocytes, cluster 3 and 4, comprise genes up and down-regulated in cyst germ cells, respectively, and correspondingly, down- or remaining at steady state levels in follicular oocytes. The expression kinetics of six top representative differentially expressed genes of each cluster are shown in Figure 4E. Enriched GO terms of the top expressed 100 genes of these clusters are shown in Figure 4F. For example, cluster 1 enriched in genes in protein folding and female gonad development, cluster 2 contained genes involved in mitochondrial metabolism and protein folding, cluster 3 and 4 were enriched in genes of ribosome biogenesis and catabolic processes. To further characterize the oocyte fate transition from cysts to follicles, genes of cluster 1 and 2 were merged and analyzed with the ClueGO algorithm. Functionally grouped term networks were obtained of genes up-regulated in follicular oocytes: genes involved in DNA conformation changes and chromatin remodeling, meiotic cell cycle regulation, mitochondrial ATP synthesis coupled electron transport and organelle fission (Figure S4A) (Supplemental Table 2). Conversely, ClueGO functional networks of genes in cluster 3 and 4 under regulated in germ cell cysts belonged to processes such as cell cycle, reproductive development, cell proliferation, oxidative stress, WNT signal and cell adhesion (Figure S4B) (Supplemental Table 3). Pseudotime trajectories of representative genes related to DNA methylation and chromatin changes, as well mitochondrial ATP synthesis present mainly in states 4 and 5, confirmed the indications of ClueGO that such processes were highly represented in oocytes engaged in follicle assembly (Figures S2D, 4G and 4H), while the expression of adhesion genes was apparently more relevant in germ cell cysts (Figure 4I).

**Fig. 4.**
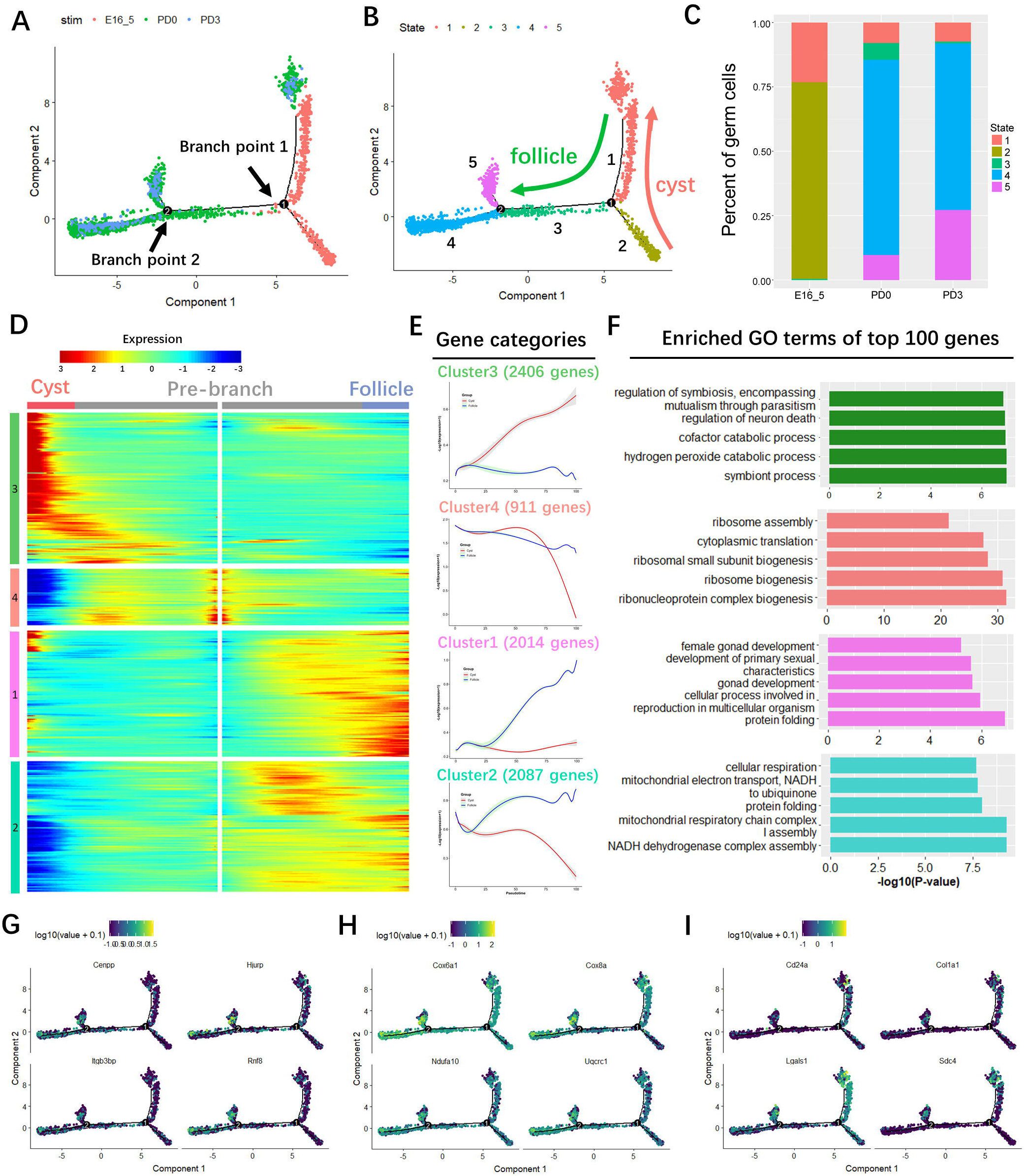
Transition of germ cells throughout distinct states from cyst to follicle stages. (A) Single-cell trajectories of germ cell subsets as a function of developmental timeline. Branch point 1 and 2 represent distinct germ cell transition states. (B) Single-cell trajectories of the five germ cell states through the pseudotime. (C) Percentage of germ cells in the five different states as a function of developmental timeline. (D) Heatmap representing the expression dynamics of four up and down expressed gene clusters at cyst and follicle stage at branch point 1 (including 1, 2 and 3 states). (E) Expression profiles of the top six differentially expressed genes in the transition from cyst to follicle in each cluster: Cluster 1: *Cacybp, Sohlh1, Id1, Lhx8, Uchl1, Figla*; cluster 2: *Mael, Taf7l, Sycp1, Fkbp6, Sycp3, Tex15*; cluster 3: *Xist, Alas2, Tmsb4x, Car2, Jun, Snca*; Cluster 4: *Rps16, Smc1b, Rps28, Rplp1, Rps23, Rpl32*. (F) Gene Ontology (GO) terms of top 100 differentially expressed genes in each cluster. (G, H and I) Expression of representative genes of chromatin remodeling (G), mitochondrial electron transport (H) and cell adhesion regulation (I) along single-cell pseudotime trajectories.

As reported above, follicular oocytes at state 3 bifurcated towards states 4 and 5 (Figure 4B). As for branch point 2, the genes of these two last states could be grouped in four distinct clusters containing up- (cluster 1, 552 genes), steady-state- (cluster 2, 332 genes) or down- regulated (cluster 3, 693 genes; cluster 4, 434 genes) genes in state 5 compared to state 4 (Figures 5A and B). Top 100 genes of cluster 1 analyzed for enriched GO and ClueGO terms (Figures 5C and 5D), belonged to a variety of metabolic processes including production of reactive oxygen species, cell cycle check points (i.e. p53) and gonad and reproductive system development. Furthermore, pseudotime trajectories and KEGG enriched analysis of genes in cluster 1 revealed among others, abundant transcripts for PI3K-Akt signaling, apoptosis and p53-asociated processes (Figures 5E and 5F and S4C). Finally, the enriched GO and KEGG terms of clusters 3 and 4 (Figures 5C and S4D), showed an ongoing decrease of biological processes such as ribosome biogenesis, oxidative phosphorylation and meiotic cell cycle as well as of pathways active in some diseases (i.e. Huntington, Parkinson and Alzheimer, NALD), and in particular, cluster 2, the steady-state generally high expression of genes in state 4 involved in gonad/female gonad development (Figures 5A and 5B and 5C).

**Fig. 5.**
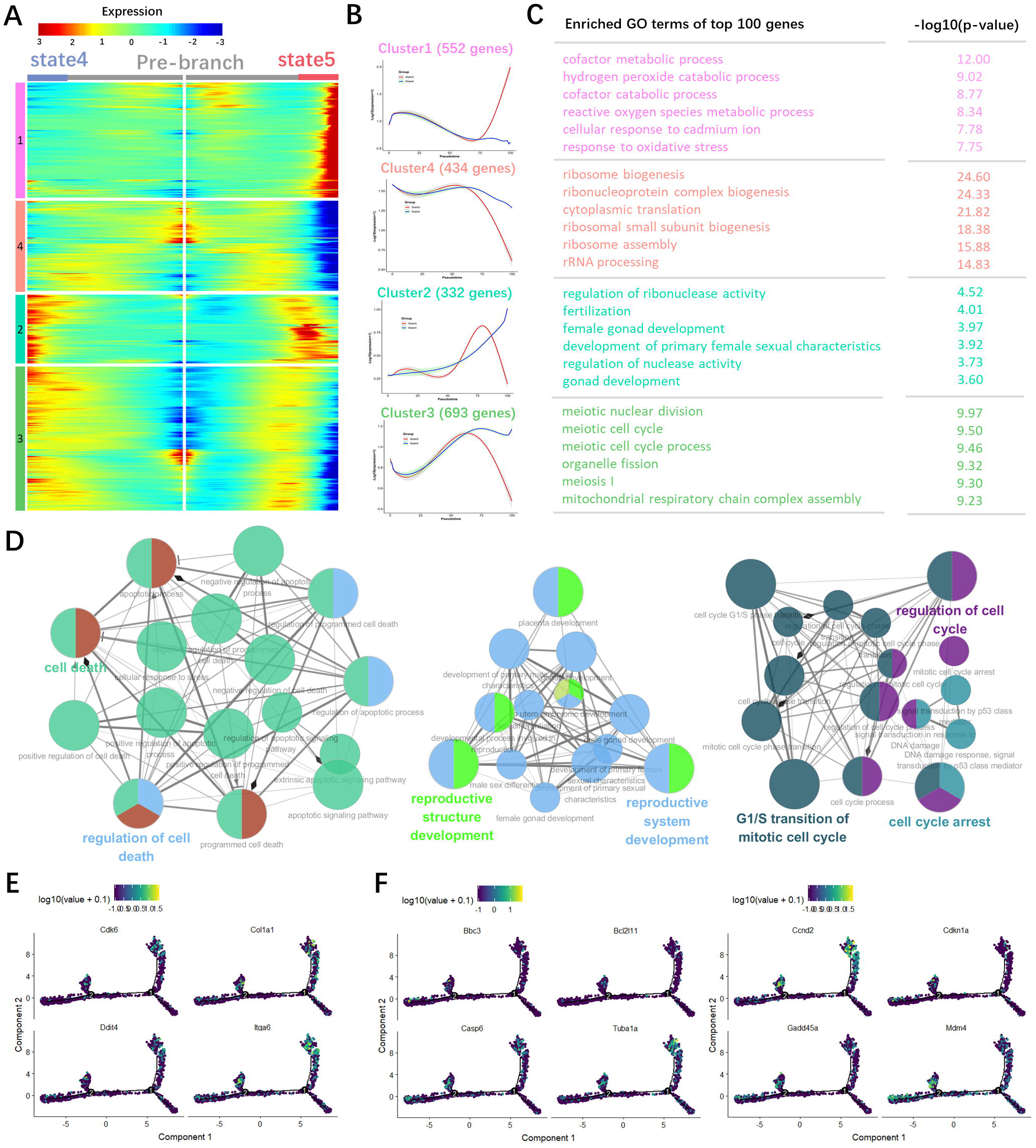
Transition of oocytes at follicle stages throughout two distinct states. (A) Heatmap showing the dynamics of gene expression in oocytes within follicles at state 4 and state 5 at branch point two. Four gene clusters of up or down differentially expressed genes of the two states. (B) Expression profiles of the six most representative genes of state 4 and 5. Cluster 1, *Xist, Hbb-bt, Hbb-bs, Hba-a1, Hba-a2, Malat1*; cluster 2, *Dppa3, Ldhb, Cd164l2, Figla, Ooep, Tmsb10*; cluster 3, *Sycp3, Mael, Taf7l, Ppia, Cirbp, Sub1*; cluster 4, *Sycp1, Rps28, Rps23, Smc1b, Rpl22l1, Rps5*. (C) GO terms of top 100 differentially expressed genes in each cluster. (D) ClueGO analysis of cluster 1 genes. (E and F) Expression of genes of PI3K (E) and apoptosis (F) pathways along single-cell pseudotime trajectories.

### Mapping oocyte specific regulon networks by SCENIC

We further investigated the regulon activity of germ cell-specific transcriptional factors (TFs) using SCENIC, an algorithm developed to deduce Global Research Network System (GRNs) and the cellular status for scRNA data (Aibar, González-Blas et al., 2017). Clustering of t-SNE were calculated based on 224 regulons activity with 9437 filtered genes with default filter parameters (Figure 6A), and the regulons density was also mapped (Figure 6B). Accordingly, the regulon activity was binarized and matched germ cell status and developmental stages (Figure 6C). As shown, four TFs, including CUX1, CREB1, GTF2F1 and NELFE, were significantly represented in all oocyte states, despite their expression being limited to specific regions. This supports these TFs as likely candidates for sustaining a germ cell specific transcription program throughout the investigated period (Figure 6D). Moreover, a series of TFs displayed a more dynamic pattern. For example, FOXO1 and BRCA1 were active mainly at the germ cell cyst stage and gradually turned off as follicular assembly occurred (Figure 6E, left). Similarly, HES1 and SOX11 seem to limit their activity to the cyst stage (Figure 6E, right), whereas HDAC2 and TAF7 expression were mainly associated to states 3 and 4, corresponding to the period of primordial follicle assembly (Figure 6F, left). Furthermore, a portion of TFs appeared to serve as a transient switch, such as STAT3 mainly at PD3 and FOXO3 in a limited period of PD0 (Figure 6F, right).

**Fig. 6.**
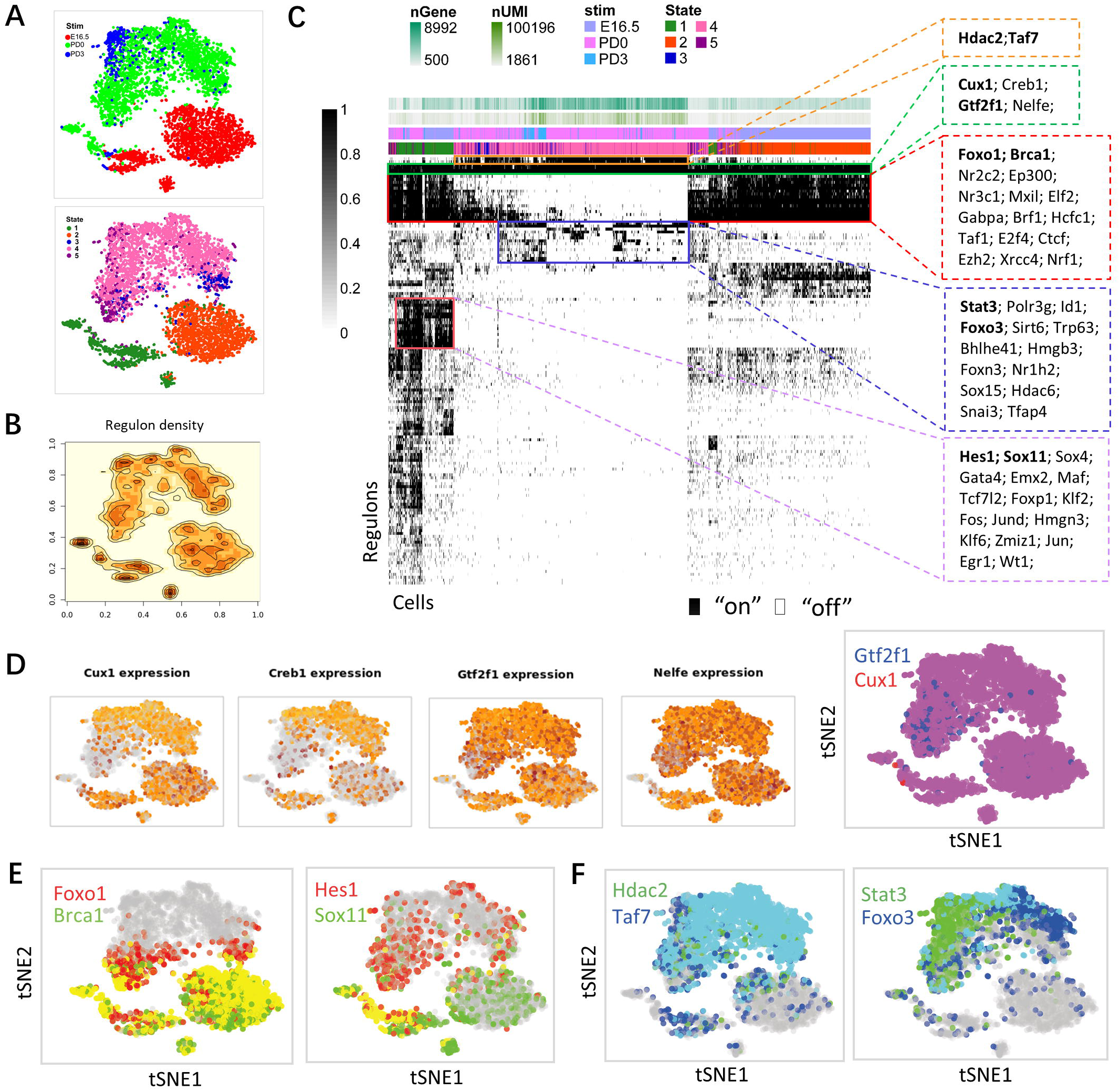
Oocyte transcription factors involved in follicle assembly. (A) t-SNE of 224 regulons based on AUC (area under the curve) values in single cell according to the developmental stages (upper) and cell states (below). The setting combination of 15 principal components (PCs), 50 perplexity (perpl) were selected for t-SNE plots. (B) t-SNE of regulons activity density. (C) Heatmap of regulon activity analyzed by SCENIC with default thresholds for binarization. “On” indicates active regulons; “Off” indicates inactive regulons. The transcription factors involved in follicle assembly were listed in the right panel. (D) t-SNE of the four conserved TFs expression and their binary regulon activity in oocytes. (E) t-SNE projection of average binary regulon activity in cyst germ cells. (F) t-SNE projection of average binary regulon activity in follicular oocytes.

### Gene expression signatures of the granulosa cell lineage

When t-SNE transcriptome of granulosa cells (Figure 7A) was plotted in function of the developmental stages, eight cell clusters were identified (Figure 7B). As note, while the expression pattern of typical granulosa cell markers such as *Wt1* and *Amhr2* were among the most conserved ones, that of *Fst* and *Kitl* varied considerably at different stages (Figure 7C). Moreover, novel marker genes within each cluster were identified, whose expressions varied according to the developmental stages (Figures S5A and S5B). For example, at PD3, *Serpine2* a gene encoding a member of the serpin family of proteins, a group of proteins that inhibit serine proteases and also expressed by granulosa cells in human adult ovary (Fan, Bialecka et al., 2019), and *Aldh1b1* and *Aldh1a1* encoding enzymes of the oxidative pathway of alcohol metabolism, reported to be associated with ovarian cancer (Marcato, Dean et al., 2011, Saw, Yang et al., 2012); at E16.5, *Cdk1c* encoding an inhibitor of several G1 cyclin/CDK complexes, a marker of juvenile granulosa cell tumors (Lamas Pinheiro, Martins et al., 2016).

**Fig. 7.**
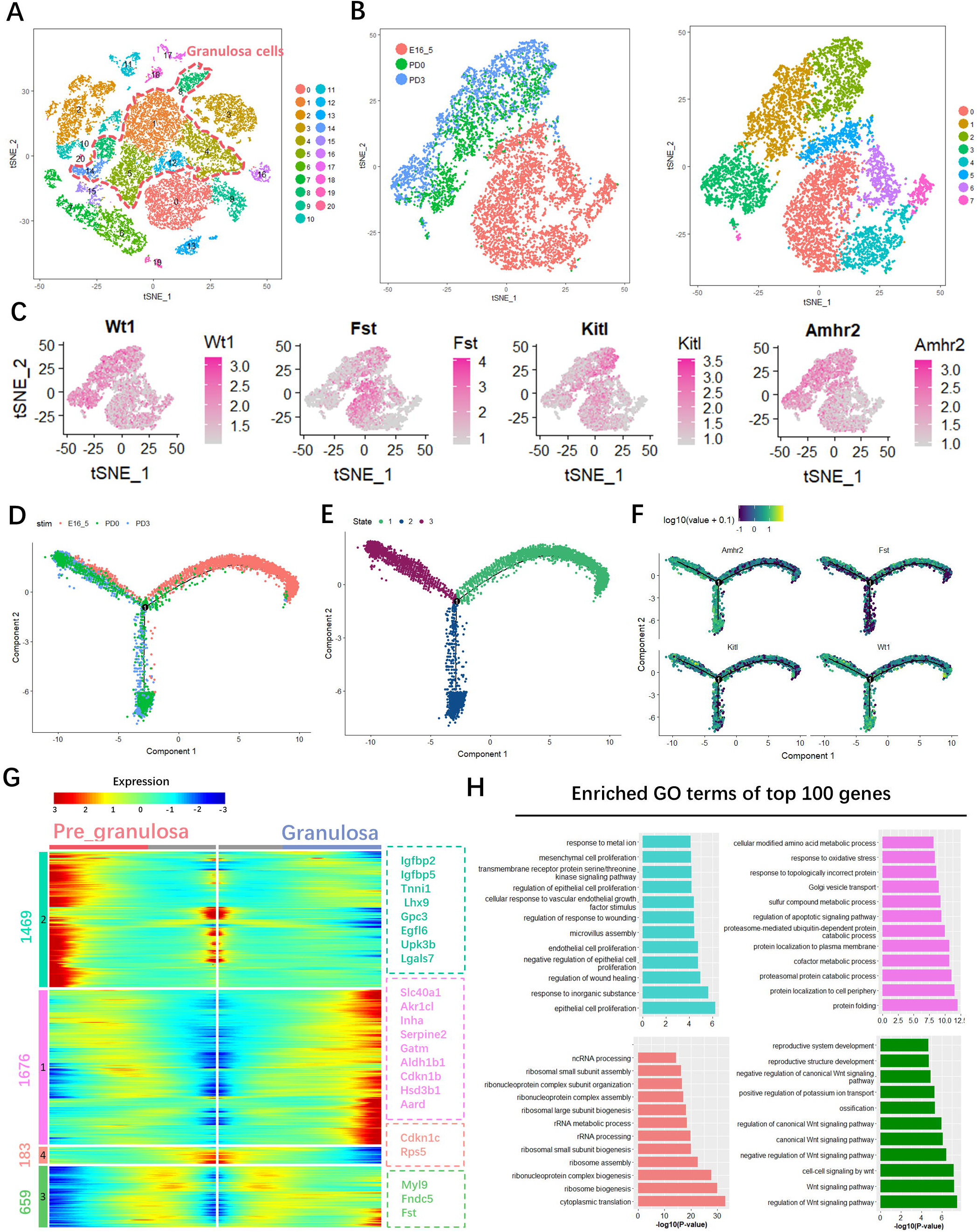
Molecular characterization of granulosa cell subsets. (A) t-SNE plot of ovarian cell clusters in which granulosa cell subsets were marked with a pink dotted line. (B) Cluster analysis of granulosa cells with t-SNE plots based on developmental timeline (left) and cell clusters (right). (C) Expression of representative genes within these clusters. (D) Single-cell trajectories of granulosa cell subsets as a function of developmental timeline. (E) Single-cell trajectories of granulosa cell clusters grouped by state. (F) Single-cell trajectories of four representative granulosa cell marker genes. (G) Heatmap of the gene expression programs from pre- to granulosa cell state. Four gene clusters represent up or down differentially expressed genes; the most variable genes are listed in the right panel. (H) Enriched GO terms of top 100 genes of each cluster.

Cell pseudotime trajectories revealed three granulosa cell states and one branch point (Figures 7D-F). Considering the notion that during mouse ovary development two waves of granulosa cell differentiation occur characterized by partly distinct gene expression (Wang et al., 2017), and on the basis of the gene expression reported in Figure S5C, we attributed state 1 to early undifferentiated pre-granulosa cells expressing the Foxl2 downstream gene *Cdkn1c* (Nicol, Grimm et al., 2018, Rosario, Araki et al., 2012). Such cells branched at state 2 to differentiate into pre-granulosa cells expressing *Lgr5, Rspo1*, or at state 3 into granulosa cells expressing *Inha* and *Hsd3b1* required for steroidogenesis (Stévant et al., 2019). For further characterization of the granulosa cell lineage, the genes of the three states were divided in four clusters according to their distinct expression patterns and a heatmap was generated (Figure 7G). Clusters 1 (1676 genes) and 3 (659 genes) were assigned to state 3 (granulosa cells), cluster 2 (1469 genes) to state 2 (pre-granulosa cells) and cluster 4 (183 genes) to state 1 (early pre-granulosa cells). On this basis, *Cdkn1c* (cluster 4) expression decreased from pre- to granulosa cells, *Igfbp2* and *Lhx9* (cluster 2), were high in pre-granulosa and declined in granulosa cells, and *Inha* and *Hsd3b1* (cluster 1) were low in pre-granulosa and high in granulosa cells (Figure 7G; Figures S5C and S5D). GO analysis uncovered that genes coding members of WNT signaling were among the enriched top 100 genes of cluster 3, containing genes low expressed in pre-granulosa cells and high expressed in granulosa cells, confirming that WNT signaling likely plays a relevant role in primordial follicle formation (Figure 7H)(Wang, Gillio-Meina et al., 2013). Cluster 1 was characterized by the expression of genes associated to biological processes such as protein folding, proteosomal processes, apoptotic pathways, and response to oxidative stress were much higher in granulosa than in pre-granulosa cells. Whereas cluster 2 was comprised of genes encoding proteins involved in mesenchymal, endothelial and epithelial proliferation that were more expressed in pre-granulosa than in granulosa cells. In line with this, when clusters 1 and 3 genes were analyzed with KEGG, terms of WNT and of FoxO pathways they were highly enriched together with that of proteosome and protein processing (i.e. folding) in the endoplasmic reticulum (Figure S5E). KEGG performed on cluster 2, showed that the most represented terms of signaling pathways in pre-granulosa cells were those of PI3K-Akt and MAPK, and to a lesser extent of Rap1 (a Ras related protein) and Hippo, while proteoglycans, tight junctions, focal adhesions, and regulation of actin cytoskeleton were among those more accounted for cellular components (Figure S5F).

### Interactions between germ cells and granulosa cells during primordial follicle assembly

Since interactions between oocytes and granulosa cells are crucial during cyst breakdown and primordial follicle assembly (Zhang et al., 2018), and throughout the entire folliculogenesis, we compared the transcriptomes of germ cells and granulosa cells together according to the developmental stages, to verify the involvement of previous discovered players and identify new cell signaling pathways driving such interactions.

Vlnplots shown in Figure S6 confirmed that at these stages NOTCH signaling, members of the TGF-beta family, Kit/Kitl system and gap junctions, are important components of the oocyte-granulosa cell interactions as previous reported (Wang et al., 2017, Zhang et al., 2018). In this regard, we found that genes encoding NOTCH ligands such as *Dll3* and *Jag1-2* were expressed by germ cells mainly at E16.5 or PD0 and PD3, respectively, while that for its receptor *Notch2* paralleled such expression in granulosa cells. The targets NOTCH gene *Hes1* showed transient expression in oocytes at PD0 and resulted in ubiquitous expression in granulosa cells at all stages, whereas *Rbpj* was more expressed by germ cells at E16.5 and PD0, and granulosa cells at PD3 (Figure S6A). TGF-beta members, *Gdf9, as* expected, was mainly found in oocytes postnatally, while the gene encoding its receptors *Bmpr1a* and *Bmpr2* were expressed by oocytes in a variable manner and continually by granulosa cells. Genes encoding the transducers SMADs and *Smad5* was expressed higher by oocytes, while *Smad3*, *Smad5* and *Smad7* were found in granulosa cells throughout the investigated period. Finally, target *Id* genes were ubiquitously expressed at all stages both by germ cells and granulosa cells (Figure S6B). Again, as expected, *Kit* was prevalently expressed by oocytes after birth and *Kitl* by granulosa cells at all stages (Figure S6C). Lastly, *Gjc1* and *Gja1* were found in oocytes and granulosa cells, respectively (Figure S6D), supporting the participation of connexin CX45 and CX43 to the oocyte-granulosa cell interaction during the entire analyzed period.

KEGG enriched analyses were used to identify bioprocesses ongoing mainly in germ cells or granulosa cells or both. For example, RNA splicing and transport followed by cell cycle regulation, meiosis and p53 signaling, were primarily found in germ cells whereas proteoglycan synthesis and focal adhesion assembly followed by WNT, FoxO and Hippo pathways, were mostly represented in granulosa cells (Figures 8A and 8B). Eight out of 44 processes found in both cell types referred to reproductive processes according to previous works (Pepling, 2012, Wang et al., 2017) (Figures 8B and 8C).

**Fig. 8.**
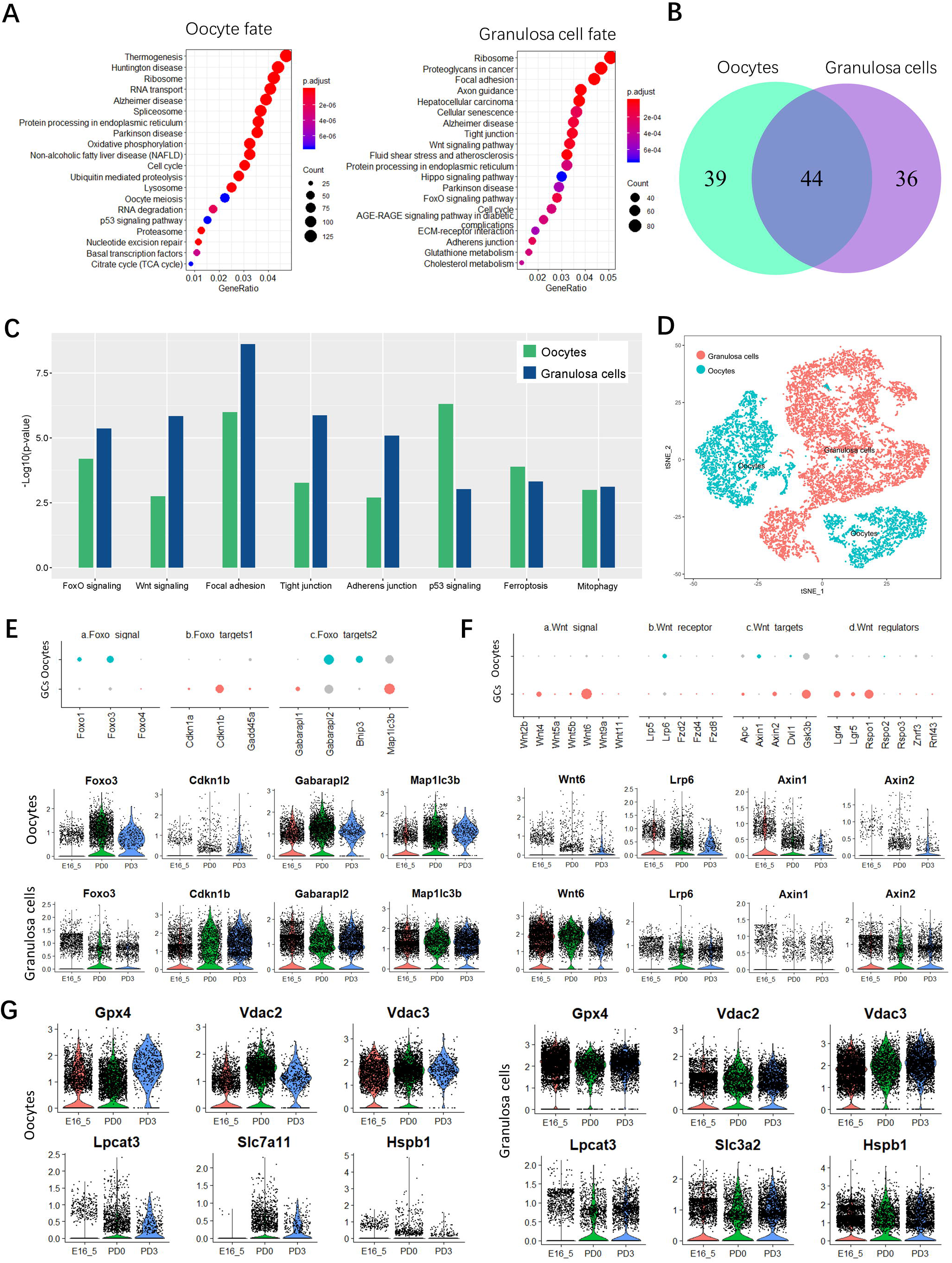
Enrichment molecular pathways of oocytes and granulosa cell subsets during follicle assembly. (A) Pathway enrichment of key genes of germ cells (7418 genes at branch point 1) and granulosa cells (3987 genes at branch point) fate transitions. (B) Vlnplots of the common and specific pathway between germ cells and granulosa cells. (C) Histogram representation of the expression level of the most representative common pathway of germ cells and granulosa cells. (D) t-SNE plot of the combined germ cell and granulosa cell subsets. (E) Dot plots of the expression of FoxO signal and its target genes in oocyte and granulosa cell clusters. (F) Dot plots (top) and Vlnplots (bottom) of expression level of WNT signal, its receptors, targets and regulators in oocyte and granulosa cell clusters. (G) Vlnplots of expression of key genes involved in ferroptosis processes in oocytes and granulosa cells.

**Fig. 9.**
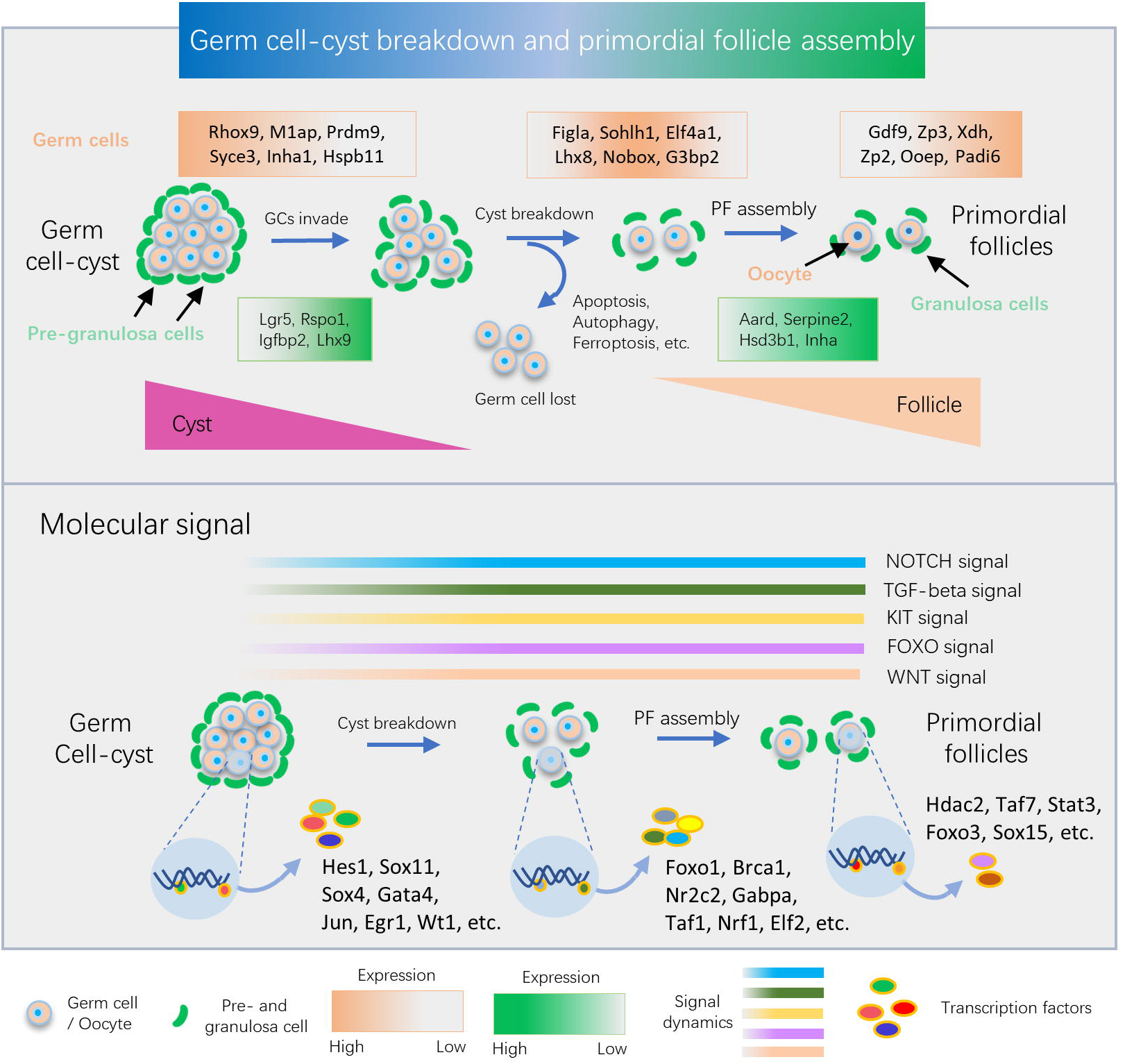
Schematic scheme of the main genetic and molecular signal dynamics detected in ovarian cells during follicle assembly. The upper panel indicates the genetic programs of germ cells and granulosa cells during germline cyst breakdown and follicle assembly, involving among others loss of about two thirds of germ cells. In the medium the activation of five main interactive pathways between germ cells and granulosa cells during these processes is represented. In the bottom, stage-specific transcription factors in oocytes are reported.

In order to characterize the crosstalk between the two cell types, germ cells and granulosa cell transcriptome clusters were combined (Figures 8D, S7A and S7B). Concerning FoxO signaling, *Foxo1* and *Foxo3* were the most common transcripts mainly expressed postnatally in oocytes and to a lesser extent in granulosa cells (Figure 8E). Among FoxO downstream genes involved in cell proliferation (target 1), *Cdkn1b* was expressed at a higher level in granulosa cells at PD3, while autophagy related genes (target 2) such as *Gabarapl1* and *2*, *Map1lc3b* and *Bnip3*, were expressed in both cell types throughout the examined stages although at different levels (Figure 8E). As for WNT signaling (Figure 8F), both *Wnt4* and *Wnt6* were expressed higher in granulosa cells than in germ cells at all stages. Transcripts for its receptor LRP6 were at similar levels in both cell types at E16.5 and PD0 and remained higher in granulosa cells at PD3. Genes encoding targets of WNT, such as *Axin1* and *Axin2* were more present in germ cells at E16.5 and at PD0, respectively, while *Axin2* and *Gsk3b* were expressed by granulosa cells at all stages. The expression of WNT6 and LRP6 were confirmed at the protein level by immunostaining (Figures S7C and S7D). Transcripts for focal adhesion proteins were more abundant in granulosa cells, those for tight junction and adherens junction proteins appeared to follow almost constantly disparate patterns (Figure S7E). Finally, nearly all genes encoding proteins involved in ferroptosis (a type of programmed cell death dependent on iron) (Galluzzi, Vitale et al., 2018, Stockwell, Angeli et al., 2017, Xie, Hou et al., 2016), were expressed at high levels in both cell types at all examined stages (Figure 8G).

## DISCUSSION

In the present study, we used scRNA-seq analyses to identify the complete transcriptome programs of the different cell types present in mouse ovaries from the end of the fetal age through the first days after birth. As reported in the Introduction, in most mammalian species, this period is crucial for ovary development and is characterized by several processes involving all ovarian cell populations. In this regard, our analyses were able to identify six distinct cell types including germ cells, pre-and granulosa cells, epithelial cells, erythrocytes, immune cells and endothelial cells that underwent dynamic changes throughout the investigated period (Figures 2A-E). These changes likely reflect the dynamics of proliferation, differentiation and death of the different cell populations that with regards to germ cells are characterized by cyst breakdown associated with extensive degeneration.

Based on the expression of known cell lineage and developmental stage-specific genes, we generated ten different gene clusters for germ cells, four at E16.5, five at PD0, and one at PD3. According to this analysis, a genetic dynamic model, including pre-, early- and late-follicle stages of germ cell development and the percent changes of each cell types at these stages were delineated (Figures 3D-F). Typical germ cell marker genes such as *Ddx4* and *Dazl*, and to a lesser extent *Dppa3*, were expressed in most of the clusters though at a higher level at early developmental stages. Genes known to be related to meiotic processes such as *Prdm9* (histone H3K4 trimethylation)(Imai et al., 2017), *Ybx2* (DNA and RNA binding)(Gu et al., 1998, Paredes et al., 2005) and *Syce3* (major component of the transverse central element of synaptonemal complexes)(Schramm, Fraune et al., 2011), showed widespread moderate or high expression except in clusters 0, 8 and 9 for *Prdm9*. On the contrary, *Stra8*, encoding a protein crucial for the beginning of meiosis (Feng, Bowles et al., 2014), was expressed at a high level in cluster 6 only as expected since its expression was reported to be limited to the early stages of meiotic prophase 1 (Tedesco, La Sala et al., 2009). With the exception of *Figla* that was expressed, at variable levels, in all clusters, all other genes such as *Lhx8*, *Nobox* and *Sohlh1* encoding transcription factors reported to be typical of follicular oocytes (Wang et al., 2017), appeared restricted to a few clusters (2, 3 and 7) assigned to the early follicle stage. Finally, *Gdf9* (encoding a growth factor of the TGF-beta family)(Wang & Roy, 2004), *Zp2* and *Zp3* (encoding zona pellucida proteins)(Dean, 2002, Zhang et al., 2018), and *Xdh* (encoding a molybdenum-containing hydroxylase involved in oxidative metabolism of purines)(Dwivedi, Arora et al., 2013, Liu, Li et al., 2019), showed an even more restricted expression limited mainly to cluster 7 corresponding to the late follicular stage.

Overall, these last results indicated the validity of our analyses and made us confident to have identified several new genes expressed by germ cells and oocytes at each of the three designated stages whose importance and role can be the focus of future studies. For example, at the pre-follicular stage, *Inca1*, *M1ap* and *Hspb11*, encoding a CDK inhibitor (Bäumer, Tickenbrock et al., 2011, Chen, Hu et al., 2008), a protein required for meiosis I progression during spermatogenesis (Arango, Li et al., 2013) and a heat shock protein that inhibits cell death through stabilization of the mitochondrial membrane (Török, Pilbat et al., 2012), respectively, are likely candidates for a role in the control of female germ cell cycle and meiosis and in their survival/apoptosis at the pre-follicular stage. At the early follicular stage, the expression of *Eif4a1*, encoding an ATP-dependent RNA helicase, a subunit of the eIF4F complex involved in cap recognition and required for mRNA binding to ribosomes (Kumar, Hellen et al., 2016), *G3bp2*, encoding an ATP- and magnesium-dependent helicase, member of nuclear RNA-binding proteins (Kobayashi, Winslow et al., 2012), *Id1*, encoding an helix-loop-helix (HLH) protein forming heterodimers with basic HLH family of transcription factors that may play a role in cell growth, senescence, and differentiation (Wang & Baker, 2015), suggest an increase of both general and specific transcription in oocytes. At the same time, the expression of *Ooep*, encoding an RNA binding protein of the subcortical maternal complex (SCMC) of the oocyte (He, Wang et al., 2018, Li, Baibakov et al., 2008) and *Padi6*, encoding a member of the peptidyl arginine deiminase family of enzymes that may play a role in cytoskeleton reorganization in the egg and in early embryo development (Esposito, Vitale et al., 2007, Yurttas, Vitale et al., 2008), support that the oocytes begin to express these proteins at the late stage of primordial follicle assembly (here indicated as late follicle). It is important to note that the experimental approach used in the present paper identifies potential candidate genes involved in the investigated events, however, gene expression changes are not always essential. Further analysis at an individual gene level is required to determine the specific function of a given gene product.

Ordering germ cells along pseudotime trajectories, we were able to reconstruct the gene dynamics involved during transition of oocytes from the pre-follicular stage (mostly germ cells in cysts) to the follicular stage (primordial follicle assembly) throughout five distinct differentiation states. This is in line with the notion that the view of defined cell types must be replaced with that of a spectrum of cell states, which are modified by the local microenvironment and produce additional layers of stochastic cell-to-cell variability. The majority of cells at pre-follicle stage belonged to states 1 and 2, while those at early and late follicle stages were in states 3, 4 and 5. Along such pseudotime trajectories, two branch points were identified. Point 1 at which the majority of cyst germ cells were at state 2 (except a small population remaining apparently at this state) considered to come from the less differentiated state 1 cells and in transit to state 3 (follicular oocytes), and point 2, at which follicular oocytes branched to the 4th and 5th states (Figures 4A and 4B and S2C). Four clusters were identified including genes likely involved in the commitment of cyst germ cells into follicular oocytes at branch point 1. Among these, the most differentially expressed were *Cacybp*, *Sohlh1*, *Id1*, *Lhx8*, *Uchl1*, *Figla* (cluster 1) and *Mael*, *Taf7l, Sycp1*, *Fkbp6*, *Sycp3*, *Tex15* (cluster2) with high and moderately increased expression, respectively, in follicular oocytes; *Xist, Alas2, Tmsb4x, Car2, Jun, Snca* (cluster 3) with high expression in cyst germ cells and down regulated in follicular oocytes, and *Rps16*, *Smc1b*, *Rps28*, *Rplp1, Rps23*, *Rpl32* (cluster 4) with low expression in cyst germ cells and up-regulated in follicular oocytes. Enriched GO and ClueGO algorithm applied to the top most expressed 100 genes and pseudotime trajectories of most representative genes involved in specific pathways of these clusters (Figures 4D-G and S4A, S4B), allow one to infer that while in cysts germ cells catabolic and adhesion processes are well represented, germ cell transition from the cyst to the follicle stage is accompanied not only, as expected, by incremental changes of processes linked to gonad/female gonad development but also by increased protein folding, mitochondrial metabolism, ribosome biogenesis, DNA conformation changes and chromatin remodeling, meiotic cell cycle regulation and organelle fission.

The same bioinformatics analyses together with enriched KEGG analysis, performed on the four cluster genes identified at the bifurcation of follicular oocytes at state 3 between state 4 and 5 at branched point 2 (Figures 5A and B), support the notion that it might be due to the early follicle dynamics characterized by the establishing in the mouse ovary of populations of dormant primordial follicles in the cortex and activated follicles in the medulla and a consistent wave of oocyte degeneration (Wang et al., 2017, Zheng et al., 2013, Zheng et al., 2014). It is possible to postulate that oocytes at state 3 and 4 belong to the dormant or activated follicles, whereas those at state 5 to degenerating oocytes. In fact, enriched GO and ClueGO of the top 100 genes in cluster 1 that were up regulated in oocytes at state 5 in comparison to state 4, and pseudotime trajectories representing genes of specific pathways, belonged to processes including production of reactive oxygen species, response to oxidative stress, cell cycle arrest (i.e. p53) and regulation of cell death/apoptosis (Figures 5C-F and S4D). On the other hand, enriched GO and KEGG terms of clusters 3 and 4 genes at state 5 showed decline of biological processes such as ribosome biogenesis, RNA metabolism, oxidative phosphorylation and meiotic regulation, mitochondrial respiratory chain complex assembly and steady-state expression of gonad/female gonad development in comparison to state 4 (Figures 5C and S4D).

Moreover, using SCENIC algorithm, we were able to establish a network of regulons that can be postulated as likely candidates for sustaining germ cell specific transcription programs throughout the investigated period continuously such as CUX1, CREB1, GTF2F1 and NELFE, or at distinct stages such as FOXO1, BRCA1, HES1 and SOX11 at the germ cell cyst stage, HDAC2 and TAF7 during the period of primordial follicle assembly and STAT3 mainly at PD3 and FOXO3 in a limited period of PD0 (Figure 6).

Bioinformatics analyses of the granulosa cell t-SNE transcriptome identified eight clusters and three states (Figure 7B). As for germ cells, the expression of genes known to be expressed by cells of the granulosa lineage during the investigated period validated these analyses showing that at the same time that some of these genes such of *Fst* and *Kitl* varied considerably at different stages. Moreover, novel genes within each cluster were identified, whose expressions changed in function with the developmental stages. Cell pseudotime trajectories revealed three granulosa cell states and one branch point (Figures 7D-E). Following the considerations reported in the results, state 1 attributed to early undifferentiated pre-granulosa cells branched at state 2 and 3 to differentiated pre-granulosa cells expressing *Lgr5, Rspo1* and granulosa cells expressing *Inha* and *Hsd3b1* required for steroidogenesis (Stévant et al., 2019). For further characterization of the granulosa cell lineage dynamics, the genes of the three states were divided in four clusters characterized by distinct expression patterns. GO analysis evidenced several biological processes, molecular functions and cellular components that increased (i.e. protein folding, proteosomal processes, apoptotic pathways, response to oxidative stress, WNT signaling) or decreased (i.e. mesenchymal, endothelial and epithelial cell proliferation) in the passage of pre-granulosa to granulosa cells. When clusters 1 and 3 genes were analyzed with KEGG, the results showed that WNT and FoxO terms as well as that of the proteosome were highly enriched (Figure S5E). KEGG on cluster 2, showed that PI3K-Akt and MAPK, and to a lesser extent Rap1 and Hippo terms, were among the most represented signaling pathways in pre-granulosa cells together to terms within proteoglycans, tight junctions, focal adhesions, regulation of actin cytoskeleton (Figure S5F).

Further, bioinformatics analyses of the germ cell and granulosa cell transcriptomes paralleled in function with the developmental stages confirmed that NOTCH signaling, members of TGF-beta family, Kit/Kitl system and gap junctions, are important components of the oocyte-granulosa cell interactions during the investigated period. It showed that ligands, receptors and signaling mediators of these system were coherently expressed mainly in germ cells, pre- and granulosa cells or both as detailed in the results. Moreover, basically in line with the finding discussed above, biological processes such as RNA splicing and transport, cell cycle regulation, meiosis and p53 signaling, were primarily found in germ cells whereas proteoglycan, cholesterol, and glutathione synthesis were more represented in granulosa cells and ribosome biogenesis in both cell types. Hippo pathways were mostly represented in granulosa cells and FoxO and WNT signaling were present in both cell types although at different developmental stages. The components of Hippo signaling in granulosa cells have been reported in several mammalian species (Kawamura, Cheng et al., 2013, Kawashima & Kawamura, 2018, Plewes, Hou et al., 2019) and disruption of the Hippo pathway has been reported to promote AKT-stimulated ovarian follicle growth (Plewes, Hou et al., 2018). Here we report its possible involvement in the early stages of granulosa cell development. As for FoxO, while a higher level of transcripts for *Foxo1* and *Foxo3* were present in post-natal oocytes, *Cdkn1b* was expressed at a higher level in granulosa cells at PD3 and autophagy related genes (*Gabarapl1, 2*, *Map1lc3b* and *Bnip3*) were expressed in both cell types throughout the examined stages although at different levels. This suggests that FoxO signaling is not only important in controlling the cell cycle (Furukawa-Hibi, Kobayashi et al., 2005) and survival of the postnatal oocytes as reported in previous studies (Pelosi, Omari et al., 2013, Sui, Luo et al., 2010), but is involved in granulosa cell proliferation and autophagy. A series of studies have identified the expression and regulation of WNT ligands and downstream WNT signaling components in developing follicles (Boyer, Goff et al., 2010, Harwood et al., 2008). The expression patterns found in our analyses for these molecules suggest that WNT pathways are among the most represented interactive signaling between the germ cells and granulosa cells during the investigated period. Interestingly, nearly all genes encoding proteins involved in ferroptosis were expressed at a high level in both cell types at all examined stages, implying this form of cell death is very active in germ cells and granulosa cells.

## MATERIALS AND METHODS

### Animals and preparation of cell suspensions

C57BL/6J strain mice were purchased from Vital River Laboratory Animal Technology Co. Ltd (Beijing, China). Animals were housed according to the national guideline and Ethical Committee of Qingdao Agricultural University. Females were mated with males in the afternoon and the presence of a vaginal plug in the morning of the following day was considered as E0.5. Embryonic and postnatal ovaries were dissected from E16.5 embryos and pups at postnatal day 0 (PD0) and 3 (PD3), respectively.

To obtain single cell populations, isolated ovaries were cut into small pieces and incubated in 0.25 % trypsin (Hyclone, Beijing, China) and collagenase (2 mg/ml, Sigma-Aldrich, C5138, Shanghai, China) for 6-8 mins at 37 °C. Tissues were disaggregated with a pipette to generate single cells, and the solution was filtered through 40 μm cell strainers (BD Falcon, 352340, USA) and washed two times with PBS containing 0.04 % BSA. Cell viability was acceptable when after staining in 0.4% Trypan Blue was above 80%.

### Single-Cell Capture and cDNA Libraries and Sequencing

Cells in 0.04 % BSA in PBS were loaded onto 10×Chromium chip and the single-cell gel beads in emulsion generated by using Single Cell 3’ Library and Gel Bead Kit V2 (10 × Genomics Inc., 120237, Pleasanton, CA, USA). Followed the manufacture instructions, the single-cell RNA-seq libraries were constructed and pair-end 150 bp sequencing was performed to produce high-quantity data on an Illumina HiSeq 2000.

### Clustering analysis with Seurat and cell trajectory construction by Monocle

The cell ranger count pipeline (version 2.2.0) were applied to the produced data to map mouse reference genome (version mm10) and process with default parameters. The data matrixes were then loaded in R (version 3.5) using Seurat package (version 2.3.4) (Butler, Hoffman et al., 2018). The Seurat object was created based on two filtering parameters of “min.cells=5” and “low.thresholds=500”, followed by the data normalize and scale, the multiple samples were combined with “RunMultiCCA” functions. Then, t-SNE plot was made by “RunTSNE” function with proper combination of “resolution” and “dims.use”. “FindAllMarkers” function was used to identify conserved marker gene in clusters with default parameter. Finally, the subpopulation of germ line or granulosa cell line was imported into Monocle (version 2.10.1) (Qiu, Mao et al., 2017) and created new dataset for monocle object, and functions of “reduceDimension” and “orderCells” were carried out to generate the cell trajectory based on pseudotime. Particularly, the state 2 cell was considered as soot state. In addition, the “BEAM” function was used to calculate the differentially expressed genes at branch point in the trajectory and gene with qval < 1e-4 showed with heatmap, expression kinetics of top six genes in each cluster were displayed with 95% confidence interval.

### The regulon activity of transcription factors with SCENIC

Following the standard pipeline, the gene expression matrix with gene names in rows and cells in columns was input to SCENIC (version 0.9.1) (Aibar et al., 2017). The genes were filtered with default parameter and finally 9,437 genes available in RcisTarget database, the mouse specific database (mm10) that default in SCENIC. The co-expressed genes for each TF was constructed with GENIE3, followed by spearman correlation between the TF and the potential target, then the “runSCENIC” procedure help to generate gene regulatory network (GRN, also termed regulons). Finally, the regulons activity was analyzed by AUCell (Area Under the Curve), whose default threshold was applied to binarize the specific regulons (on or off). The default of tSNE was settled with PCs (principal component) for 15 and perpl (perplexity) for 50, meanwhile the cell states were mapped with specific regulons, and AUC and TF expression were projected onto t-SNEs. Enrichment analysis of gene ontology (GO) and Kyoto encyclopedia of genes and genomes ClusterProfile (Yu, Wang et al., 2012), a R package in Bioconductor, was applied to detect the gene related biological process and signaling pathways with threshold value of “pvalueCutoff = 0.05”, and top terms was displayed (Figures 4F, 5C, 7H, 8A, S4C, S4D, S5E and S5F). ClueGO (V2.3.5), that an implanted plug-in Cytoscape (V 3.4.0), was used to uncover the functional units that related reproductive development (Figures S5D, S4A and S4B)(Bindea, Mlecnik et al., 2009).

### Immunohistochemistry

Immunohistochemistry was performed as previously described (Wang, Yu et al., 2018). Ovarian samples were processed for paraffin inclusion after fixing with paraformaldehyde (Solaibio, P1110, China) and following a standard dehydration procedure. Sample sections of 5 μ thickness were blocked at room temperature for 30 min after gradient rehydration and antigen retrieval, and incubated with the germ cell specific MVH antibody (Abcam, ab13840, USA) overnight at 4 °C. Followed with three wash in TBS, the sections were incubated with secondary antibodies of FITC-labeled goat anti-rabbit IgG (H+L) (Beyotime, A0562, China) for half an hour at 37 °C. Eventually, after TBS washes three times, slides were counterstained with propidium iodide (PI, Sangon Biotech, E607306, China) for 3 min and sealed, the pictures were taken with a fluorescence microscope (Olympus, BX51, Japan). Double immunostaining was performed with MVH antibody from mouse (Abcam, ab27591), rabbit HSPB11 (15732-1-AP, Proteintech, USA), G3BP2 (A6026, ABclonal, USA), XDH (55156-1-AP, Proteintech), WNT6 (24201-1-AP, Proteintech) and LRP6 (A13324, ABclonal) antibodies, followed by secondary antibodies of CY3-labeled goat anti-mouse IgG (H+L) (Beyotime, A0521) and goat anti-rabbit IgG H&L (Alexa Fluor® 488, Abcam, ab150077); Hoechst 33342 or propidium iodide (Beyotime, C1022) were used for nucleus counterstaining

### Statistical analysis

Results data as mean ± SD were obtained from three independent experiment. The statistical analysis was performed with GraphPad Prism software (version 5.0), and the significant difference was determined with two-tailed student’s unpaired *t*-Test. Significant and highly significant difference were set at *P <* 0.05 and *P <* 0.01, respectively.

## ACKNOWLEDGEMENTS

This work was supported by National Key Research and Development Program of China (2018YFC1003400), National Nature Science Foundation (31671554 and 31970788) and Taishan Scholar Construction Foundation of Shandong Province.

## Author contributions

J.J.W., W.G., Q.Y. Z., J.C.L., X.W.S., W. X. L. and L.L. conducted the experiments; J.J.W. analyzed the data; J.J.W., W.S., C.Z.L., P.W.D. and M.D.F. wrote the manuscript; W.S. designed the experiments. All authors reviewed and revised the manuscript.

## Data Availability

The accession number of ovarian single-cell RNA sequencing data reported in this paper is NCBI GEO: GSE134339.

## Supplemental Tables

Supplemental Table 1: Top 50 marker genes of cell clusters in combined data (CSV 71kb);

Supplemental Table 2: ClueGO Result of Germ cell at Branch1 of genes in cluster 1 and 2 (XLS 1058 kb)

Supplemental Table 3: ClueGO Result of Germ cell at Branch1 of genes in cluster 3 and 4 (XLS 2640 kb).

## Supplemental Figures

**Fig. S1.** (A-C) Cluster analysis of *Ddx4* and *Dazl* gene expression at E16.5(A), PD 0 (B) and PD 3 (C). (D) Canonical correlation analysis (CCA) of all ovarian cells according to the three developmental stages. (E) Vlnplots of all ovarian cells at three developmental stages under CC1 and CC2 condition. (F) Conserved genes in CC1 and CC2 integrated analyses. (G) Heatmap of the top 10 conserved marker genes in clusters after integrated analysis.

**Fig. S2.** (A) Feature plots of known germline marker. (B) Vlnplots indicating the expression profile of known marker genes along the developmental stages. (C) Cell trajectories of marker gene expression in germ cell clusters of pre-, earlyx- and late-follicle stages. (D) Cell trajectories and Vlnplots of representative genes involved in DNA methylation changes.

**Fig. S3.** (A) Representative images of the HSPB11 immunohistochemistry staining of ovary sections from E16.5 embryos, and PD0 and PD3 pups. (B) Representative images of the G3BP2 immunohistochemistry of ovary sections from E16.5 embryos, and PD0 and PD3 pups. (C) Representative images of the XDH immunohistochemistry staining of ovary sections from E16.5 embryos, and PD0 and PD3 pups.

**Fig. S4.** (A) ClueGO terms of gene clusters 1 and 2 up-regulated in follicular oocytes. (B) ClueGO terms of gene clusters 3 and 4 unregulated in cyst germ cells. The different nodes are the enriched pathways, lines in different nodes represent common genes and colors refer to the classification of enriched groups. (C) Enrichment of KEGG pathway of gene cluster 1 at branch point 2. (D) Enrichment of KEGG pathway of gene clusters 3 and 4 at branch point 2.

**Fig. S5.** (A) Heatmap of the high variable gene (top 5) in granulosa cell clusters. (C) Vlnplots of these genes in granulosa cell clusters according to the developmental stages. (D) Expression profiles of the six most variable genes of granulosa cell in gene cluster 1, 2, 3 and 4. (E) Enrichment of KEGG pathway of gene clusters 1 and 3. (F) Enrichment of KEGG pathway of gene cluster 2.

**Fig. S6.** (A) Vnlplots of the expression of NOTCH signal ligands, receptors and targets in germ cells and granulosa cells. (B) Vnlplots of the expression of TGF-beta signal ligands, receptors, effectors and targets in germ cells and granulosa cells. (C) Vnlplots of the expression of Kit and Kitl in germ cells and granulosa cells. (D) Vnlplots of the expression of connexin genes of gap junction in germ cells and granulosa cells.

**Fig. S7.** (A) t-SNE of the combined germ cells and granulosa cells subsets at the three developmental stages. (B) Feature plots of germ cells and granulosa cells identified with specific marker genes. (C) Representative images of WNT6 immunohistochemistry of ovary sections from E16.5 embryos, and PD0 and PD3 pups. (D) Representative images of LRP6 immunohistochemistry of ovary sections from E16.5 embryos, and PD0 and PD3 pups. (E) Dot plots and Vnlplots of focal adhesion, tight and adherens junction genes in germ cells and granulosa cells.

